# Agronomic characterisation of a common buckwheat germplasm collection unlocks resources for genomics-assisted ideotype breeding

**DOI:** 10.64898/2026.10.09.757874

**Authors:** Fabian Hess, Lukas Kronenberg, Gabriele Rimat, Heidi Heuberger, Matthias Heinrich Herrmann, Vera Benyr, Steven Yates, Bruno Studer

## Abstract

The mechanisation of agriculture has achieved major increases in labour productivity but has also led to an erosion of agrobiodiversity. Neglected and underutilized crops such as common buckwheat (*Fagopyrum esculentum* Moench) offer the potential to diversify food production systems but require major genetic improvement to compete with established crops. Although target traits for breeding such as more homogeneous ripening, reduced height, larger seeds and higher and more stable yields are well known, large-scale and field-based phenotyping data of diverse common buckwheat accessions across multiple environments are still scarce. Here, we tested the agronomic performance of a common buckwheat germplasm collection consisting of 170 accessions of diverse origin. The collection was tested at seven year-sites, and 19 agronomic traits related to phenology, growth habit and yield were assessed. Most traits showed high repeatability (0.31 < R^2^ < 0.98) and heritability (0.45 < H^2^ < 0.97), indicating genetic control of these traits. Coefficients of variation ranged between 6.19% and 46.15%, showing the high phenotypic diversity within the collection. By combining multiple traits, distinct ideotypes for different roles in the crop rotation were investigated, and accessions with promising agronomic profiles identified. Genome-wide association studies revealed marker-trait associations for 18 traits, allowing the identification of putative candidate genes influencing these traits such as a putative *GRF-INTERACTING FACTOR* associated with flowering time and biomass production. Our results show that the collection offers vast phenotypic diversity that can be exploited for the development of improved varieties tailored to meet specific agronomic purposes. The efficient utilization of these genetic resources should be supported by the application of genomics-based breeding approaches. The development of genetic markers and search for putative candidate genes mark important steps towards the establishment of efficient breeding schemes. Thus, these genomic resources provide a foundation for accelerating the genetic improvement of common buckwheat and developing agronomically competitive varieties.

## 2 Introduction

Common buckwheat (*Fagopyrum esculentum* Moench) is a valuable addition to most crop rotations due to its short growing season, resistance to pests and diseases and positive effects on soil fertility (Campbell, 1997; Farooq *et al*., 2016). Moreover, common buckwheat provides nectar and pollen for insects, fostering ecosystem functioning, and offers a gluten-free alternative to cereals (Joshi *et al*., 2020) with a favourable protein composition (Zeller and Hsam, 2004; Škrabanja and Kreft, 2016). However, in Europe, the once widely cultivated crop (Aufhammer *et al*., 1995; Miedaner and Longin, 2017) has largely been replaced by high-yielding cereals and other crops such as potato (*Solanum tuberosum* L.; Zeller and Hsam, 2004). As a consequence, little effort and resources have been directed towards common buckwheat improvement (Joshi *et al*., 2019) and main agronomic challenges such as low and unstable yields as well as uneven ripening remain largely unresolved (Jacquemart *et al*., 2012).

In addition to the lack of breeding activities, information on the agronomic performance of diverse common buckwheat germplasm across Europe remains scarce. Although in the 1980s and 1990s, several large-scale field trials were conducted in Nepal, China, and Japan (Campbell, 1997), and more recently in India (Senthilkumaran *et al*., 2008; Naik *et al*., 2022), these investigations do not reflect the environmental conditions of Western Europe. Moreover, they mainly focused on local landraces, which do not represent accessions cultivated in Europe. While there have been some investigations into the agronomic performance of common buckwheat under European field conditions in recent years, these investigations were limited to small numbers of accessions (n <=33, Siracusa *et al*., 2017; Strahm *et al*., 2019; Grimes *et al*., 2025). Thus, the results do not represent the available diversity of common buckwheat germplasm and offer only limited opportunities to employ methods of quantitative genetics.

Besides the availability of reliable data, a clear understanding of the target ideotype is crucial for successful breeding. Moving away from focusing on single traits like yield and towards a concept of ideotypes can help to gain a more comprehensive understanding of breeding targets (Donald, 1968). A “cereal-like” common buckwheat ideotype for seed production has been described in the literature, combining reduced hight with determinate growth habit and erect positioning of the leaves (Amelin *et al*., 2020). However, more detailed descriptions of ideotypes of common buckwheat are required similar to the ideotype for Tartary buckwheat (*Fagopyrum tataricum* Gaertn) proposed by Li *et al*. (2026). Moreover, differentiated ideotypes related to specific roles of common buckwheat in the agricultural production system should be defined such as a main or second crop for grain production (Radics and Mikóházi, 2010), fodder crop (Amelchanka *et al*., 2010), cover crop (Taranenko *et al*., 2016) or melliferous flower crop (Halbrecq *et al*., 2005).

The breeding for accessions that fit specific ideotype requirements is a necessary step to make the crop economically attractive to farmers and increase utilization. Such a development will also become increasingly necessary to meet the goals of agricultural policies that focus on reducing the environmental impact of food systems such as the European Green Deal (Sotte and Vergamini, 2025). Currently, common buckwheat breeding is limited to population breeding approaches, making the fixation of desirable alleles in the breeding population complicated and slow (Aii *et al*., 1998). The implementation of genomic tools to speed up breeding has been recognized as a necessary step to boost agronomic performance of orphan crops in general (Varshney *et al*., 2012; Nature Communications Editorial Staff, 2024) and common buckwheat in particular (Yabe *et al*., 2018; Joshi *et al*., 2019; Nay *et al*., 2020), but their practical application is still lacking. First steps towards conducting genome-wide association studies (GWAS) with common buckwheat have been taken using Himalayan landraces of common and Tartary buckwheat (Naik *et al*., 2024; Rashid *et al*., 2026). However, only 68 accessions of common buckwheat were included and the genotypic data was restricted to biallelic single nucleotide polymorphisms (SNPs), which do not adequately represent the genetic makeup of accessions of outcrossing crops such as common buckwheat (Byrne *et al*., 2013; Nay *et al*., 2020).

Meanwhile, resources and methods to advance common buckwheat breeding towards genomics-assisted approaches have become increasingly available. Multiple high quality reference genome assemblies for common buckwheat have been published (Fawcett *et al*., 2023; Lin *et al*., 2023; Hess *et al*., 2026*a*) and genome-wide allele frequency fingerprints (GWAFFs) based on double digest restriction-site associated DNA sequencing have been established as an adequate genotyping method to capture allelic diversity among accessions of common buckwheat (Nay *et al*., 2020; Hess *et al*., 2026*b*). Recently, a large population comprising 170 accessions of common buckwheat from diverse global origins has become available (ETH Zurich/FOAG (Federal Office for Agriculture) collection; Hess *et al*., 2026*b*).

The main objective of this study was to assess the performance of the accessions in the ETH Zurich/FOAG collection (henceforth referred to as the collection) for key agronomic traits related to phenology, growth habit and yield across multiple environments. Based on these data, we aimed at exploring the phenotypic diversity across the collection, suggesting ideotypes for different roles within agricultural systems and providing information for selecting future breeding material. Furthermore, we explore the underlying genomic basis of the investigated traits through GWAS, to foster genomics-assisted breeding in common buckwheat.

## 3 Materials and methods

### 3.1 Plant material, experimental design and genotyping

Field trials were conducted at three and four locations in 2023 and 2024, respectively, totalling seven year-sites located in Germany and Switzerland (Supplementary Figure S1G, Supplementary Table 1). Different environmental conditions were recorded for the year-sites in terms of weather (Supplementary Figure S1A-D and F), soil type and crop rotation (Supplementary Table S1). The complete panel of common buckwheat investigated in this study comprises 170 accessions of diverse origin, including traditional and commercial accessions, as well as nine self-compatible accessions (Matsui *et al*., 2003), and eleven accessions with determinate growth habit (Fesenko *et al*., 2018). The collection was described in detail in Hess *et al*. (2026*b*). Not all accessions could be included in each trial and the number of replications varied depending on available space and resources at each year-site (Table 1). Four trials were designed as randomized complete block designs with two (JKI23, LfL24) and three (ETH23, ETH24) replications while LfL23 only consisted of one replication. At JKI24 and HOH24 genotypes were grouped into tree ripening groups and separate randomized complete block experiments with two complete blocks were conducted for each group (Table 1). Check accessions were included in all trials to better account for spatial variability within fields. The number of check accessions and their replication varied depending on the number of experimental plots available at each year-site (Table 1). The trials were sown in 3 m^2^ plots with a sowing density of 120 viable seeds m^-2^ between 15^th^ of May and 26^th^ of June, depending on weather and soil conditions at the respective year-site.

**Table 1:** Description of trial year-sites. Reps. = Replications, Acc. = Accessions, Check acc. = number of check accessions that are replicated multiple times within replications, Harvesting groups = Harvesting groups in trails where plots are grouped in three main blocks with different harvesting dates (early, middle, late).

| Year-site | Coordinates | Plots | Reps. | Acc. | Check acc. | Check plots | Sowing date | Harvesting groups |
| --- | --- | --- | --- | --- | --- | --- | --- | --- |
| ETH23 | 47.440150, 8.689486 | 594 | 3 | 165 | 4 | 96 | 31/05/2023 | No |
| ETH24 | 47.443583, 8.681289 | 567 | 3 | 166 | 4 | 60 | 15/05/2024 | No |
| JKI23 | 54.071746, 12.315323 | 360 | 2 | 167 | 1 | 24 | 09/06/2023 | No |
| JKI24 | 54.071601, 12.312034 | 360 | 2 | 168 | 1 | 24 | 22/05/2024 | Yes |
| LfL23 | 48.700431, 11.530585 | 216 | 1 | 161 | 7 | 54 | 24/05/2023 | No |
| LfL24 | 48.700431, 11.530585 | 360 | 2 | 169 | 1 | 24 | 26/06/2024 | No |
| HOH24 | 48.523222, 7.871169 | 360 | 2 | 163 | 2 | 34 | 07/06/2024 | Yes |

The genetic characterization of the common buckwheat accessions based on GWAFFs, as well as the filtering procedure applied, have been described in Hess *et al*. (2026*b*). In short, 100 leaves per accession were sampled from different plants in the trials ETH23 and ETH24 to form two pooled samples per accession. From these samples, DNA was extracted to construct double digest restriction-site associated DNA libraries for sequencing. The resulting reads were mapped to haplome 2 of the reference genome Tuka (Hess *et al*., 2026*a*) and the two resulting BAM files of each accession were concatenated to a single file which was used for subsequent SNP calling and the calculation of allele frequencies. This resulted in a matrix of 52,182 SNP allele frequencies after filtering for a minor allele frequency of 0.05. Missing values were imputed through linkage-informed k-nearest neighbour-derived weighted mean allele frequency prediction using imputef (v.1.1.1; Paril *et al*., 2025).

### 3.2 Collection of phenotypic data

The collection was assessed for 19 agronomically relevant traits related to establishment and crop stand development, phenological progress and yield (Table 2). Emergence was recorded once seedlings had appeared in all plots (around 14 days after sowing, DAS) by randomly placing a one-meter scale at three locations within each plot and counting all seedling growing within these three meters. Canopy cover was assessed at two distinct timepoints, once when on average around 50% of each plot was covered (around 28 DAS, Cover) and once when 75% of each plot was covered (around 35 DAS, Cover late). Canopy cover was measured using digital RGB images taken by smartphones, or LITERAL (HIPHEN, Avignon, France), depending on the experiment, and processed using the Canopeo app (Patrignani and Ochsner, 2015). Early vigour was visually assessed by grading each plot with a score from 1 (low) to 9 (high) on the same date as Cover was measured. Intermediate height (around 80 to 90% of final height reached) and Final height were measured using a tape measure attached to a stick placed in the middle of each plot to assess the average canopy height. At the year-sites ETH23 and ETH24, BBCH stages (Lancashire *et al*., 1991) were recorded twice per week during the entire growing season and development stages First flower (DAS), First flower TINF (Terminal inflorescence) (DAS), First green fruits (DAS) and First brown fruit (DAS) were extracted by fitting a monotonically increasing *p*-spline to the data of each plot. The phenological development was also assessed at JKI23 (only First flower (DAS) was graded) and LfL24, by scoring plots during the respective time periods for the respective BBCH stages. Branching and Lodging were scored with grades 1 (low) to 9 (high) at an appropriate time during the field season depending on the conditions at each year-site.

**Table 2:** Description of the 1S agronomic traits assessed in the field trails, including the unit of the trait, a short description and the year-sites where the trait was assessed with the number of year-sites in brackets. The grades used for evaluating Branching, Early vigour and Lodging all range from 1 (low) to S (hight). The abbreviation DAS refers to “days after sowing”.

| Trait name | Unit | Description | Year-sites (number) |
| --- | --- | --- | --- |
| Biomass | kg plot<sup>-1</sup> | Dry, above-ground biomass without seeds | ETH23, ETH24 (2) |
| Branching | grade | Degree of primary and secondary branching 1 (low) to 9 (high) | HOH24, LfL23 (2) |
| First brown fruits (DAS) | days | Brown seeds visible in around 50% of plants | ETH23, ETH24, LfL24 (3) |
| Cover | % | Percent of plot covered with leaves (around 21 DAS) | ETH23, ETH24, JKI23, JKI24, LfL23, LfL24 (6) |
| Cover late | % | Percent of plot covered with leaves (> 21 DAS) | ETH23, ETH24 (2) |
| Early vigour | grade | Vigour of early development (“cover × height”) 1 (low) to 9 (high) | ETH23, ETH24, LfL23, LfL24 (4) |
| Emergence | count | Number of seedlings on 3 × 1 m | ETH23, HOH24, LfL23, LfL24 (4) |
| Final height | cm | Canopy height at harvest | ETH23, ETH24, HOH24, JKI23, JKI24, LfL23, LfL24 (7) |
| First flower (DAS) | days | Flower visible in around 50% of plants | ETH23, ETH24, JKI23, LfL24 (4) |
| First flower TINF (DAS) | days | Flower visible in terminal flower cluster around 50% of plants | ETH23, ETH24, LfL24 (3) |
| First green fruits (DAS) | days | Green seeds visible in around 50% of plants | ETH23, ETH24, LfL24 (3) |
| Harvest (DAS) | days | Date of harvest | ETH23, ETH24, LfL23, LfL24 (4) |
| Intermediate height | cm | Canopy height during development | ETH24, LfL23, LfL24 (3) |
| Lodging | grade | Severity of lodging in plots assessed when feasible 1 (low) to 9 (high) | ETH23, ETH24, HOH24, LfL23, LfL24 (5) |
| Proportion of ripe fruits | % | Percent of ripe fruit (based on color) in the field assessed shortly before harvest | JKI23, JKI24 (2) |
| Seed length | mm | Average length (longer side) of seeds | ETH23, ETH24, JKI24 (3) |
| Seed width | mm | Average width (shorter side) of seeds | ETH23, ETH24, JKI24 (3) |
| TGW | g | Thousand grain weight | ETH24, JKI24 (2) |
| Yield | kg plot<sup>-1</sup> | Seed yield | ETH23, ETH24, HOH24_early, HOH24_mid, JKI23, JKI24_early, LfL23, LfL24 (8) |

Plots were either directly harvested with a plot combine or first cut by hand and manually fed to a plot combine positioned at the side of the field (Supplementary Table S1). The threshed seeds were dried and threshed again in a stationary lab thresher and weighed. Biomass was measured during seed harvest by weighing the biomass (BM_fresh_) ejected by the combine harvester after threshing. From each plot, a sample of fresh biomass (around 100g) was collected, weighed (S_fresh_) and dried at 70° C to stable weight (about two days), and weighed again (S_dry_) to calculate dry biomass (BM_dry_) according to:

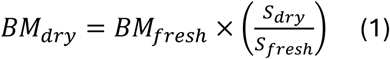

The Proportion of ripe fruits, Seed length, Seed weight and Thousand grain weight (TGW) were assessed after the seed was cleaned. While the Proportion of ripe fruits was graded by assessing the proportion of ripe fruits in the field shortly before harvest, a MARViN ProLine II (Marvitech, Wittenburg, Germany) was used to measure seed length, width and TGW.

### 3.3 Statistical analysis

All statistical analyses were performed in R (v4.4.0) unless stated otherwise.

#### 3.3.1 Spatial correction and repeatability

The traits assessed on plot level were analysed in a two-stage linear mixed model approach as described in Roth *et al*., (2021) and Roth *et al*., (2023). In the first stage, the R package SpATS (v.1.0.19) (Rodríguez-Álvarez *et al*., 2018) was used to calculate adjusted genotype means for each year-site. The model parameters were specified as:

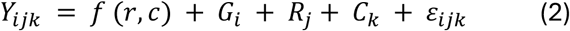

where *Y_ijk_* is the trait value of the plot level of the *i*^th^ genotype, *j*^th^ year-site and of the *k*^th^replication, *f* (*r*, *c*) is a smooth bivariate surface defined over numeric row and range positions, *G_i_* is the effect of the *i*th genotype while *R_j_* and *C_k_* are the effects of the *j*th row and *k*th range coded as factors, respectively, and s*_ijk_* refers to the residual. To calculate repeatability (Oakey *et al*., 2006) on the year-site level, *G_i_* in Equation 3 was set as random while for the calculation of best linear unbiased estimators (BLUEs) the term was set as fixed. The diagonal of the inverse of the variance-covariance matrix of the BLUEs was extracted from the model and used to weight the residuals in the next stage.

For the calculation of genotype means for yield at year-sites where accessions were grouped into three different ripening groups (Supplementary Table S1), each ripening group was analysed separately. For HOH24, only the ripening groups early and mid were used for further analysis, while for JKI24 only the group early was included. For all other traits, all three groups were analysed in a single model with ripening group being added as a covariable.

#### 3.3.2 Genotype means across year-sites and heritability

To calculate genotype means across year-sites, the R package ASReml-R (v.4.1.0.176) was used to specify the model:

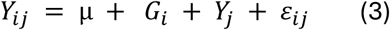

where *Y_ij_* are the BLUEs from the first stage, µ the global intersect, *G_i_* the genotype effect and *Y_j_* the year-site effect, and s*_ij_* the weighted residuals. The *G_i_* term in Equation 4 was set as random to calculate heritabilities (Cullis *et al*., 2006), while it was specified as fixed for the calculation of BLUEs across years-sites.

#### 3.3.3 Yield stability

To assess yield stability across environments, Equation 4 was extended to a factor analytic model with the *GY _ij_* term specified in ASReml-R as fa(location, 3):id(gen). Based on this model, the type B genetic correlation between year-sites was calculated, and the loadings of the three factors extracted using ASRtriala (v.1.0.2) (Gezan *et al*., 2022). Moreover, yields at all locations were predicted to calculate Wricke’s ecovalence (Wricke, 1962) for all accessions according to:

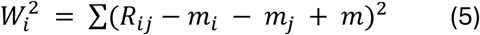

where *W*_*i*_^2^ represents Wricke’s ecovalence, *R*_*ij*_ represents the yield (BLUE) from the first stage (per year-site), *m*_*i*_ is the genotype mean yield across year-sites, *m*_*j*_ is the year-site mean yield genotypes and *m_j_* represents the grand mean of the yield.

#### 3.3.4 Phenotypic correlations between traits

All phenotypic correlations between traits were calculated based on BLUEs over year-sites. Two-sided *p*-values were calculated for each pair of traits and adjusted for multiple testing based on Benjamini-Hochberg correction, and three different significance levels were defined: α < 0.05, α < 0.01 and α < 0.001. The function hclust from the R package stats (v.4.5.1) was used for hierarchical clustering to order traits in figures and define groups of similar traits.

#### 3.3.5 Genome-wide association studies and their interpretation

The R package GAPIT (v.3.5.0; Wang and Zhang, 2021) was used for GWAS on all traits and available year-sites, including means across year-sites. The phenotypic data was introduced as BLUEs, and the genotypes were supplied as an imputed allele-frequency matrix comprising 52,182 SNP allele frequencies. Four methods implemented in GAPIT were used: BLINK (Huang *et al*., 2019), FarmCPU (Liu *et al*., 2016), mixed linear model (MLM; Yu *et al*., 2006) and multi-locus mixed model (MLMM; Segura *et al*., 2012). To account for population structure within the collection, the first three principal components of the genomic matrix were used as covariates, and the relatedness between accessions was modelled using the genomic kinship matrix proposed by (VanRaden, 2007). For each run, the significance of marker-trait associations (MTA) was assessed using the Benjamini-Hochberg FDR with significant defined as α <0.05.

To further investigate the genetic basis of the observed traits, the positions of significant MTAs on the reference genome Tuka were extracted and all genes annotated in Tuka within 50 kb of each marker were retrieved. The predicted protein sequences for these genes were used as input for EggNOG-Mapper (Cantalapiedra *et al*., 2021) with default parameters. This tool uses precomputed orthologous groups and phylogenies from the EggNOG (v.5.0.0) database (Huerta-Cepas *et al*., 2019) to transfer functional information to the queried sequences. Additionally, for all significant MTA detected within a gene, a linear model was used to model the allelic effect, where the *r*^2^ coefficient was used to describe the percentage of phenotypic variation explained.

## 4 Results

### 4.1 Phenotypic distribution and heritability of traits

A total of 19 traits were phenotyped at a minimum of two years-sites for all accessions in the collection (Table 2). Mean, maximum and minimum values, coefficients of variation and across year-sites heritabilities are summarized in Table 3, while repeatabilities for the respective year-sites are presented in Supplementary Tables S2 to S11. Most traits showed strong genotypic variation as evidenced by high observed heritabilities (H^2^ > 0.8), with the exceptions of Intermediate height, Lodging, Branching and Emergence, which showed moderate heritabilities of 0.72, 0.61, 0.51 and 0.45, respectively (Table 3). A similar pattern could be observed for within year-site repeatabilities (Supplementary Tables S2 – S11). However, the repeatabilities at year-sites with lower numbers of replications tended to be lower (especially for LfL23 with only one replication and repeatabilities calculated based on check accessions). The coefficients of variation for the across year-site BLUEs ranged from 6.19% (Seed length) to 20.56% (Lodging) for most traits (Table 3). However, Biomass, Yield and Proportion of ripe fruits showed considerably higher coefficients of variation with values reaching 36.32%, 43.17% and 46.15%, respectively. In absolute values, BLUEs across year-sites for traits with high variability such as Yield and Biomass ranged from 0.03 to 0.87 kg plot^-1^ and 2.11 to 6.54 kg plot^-1^, respectively, while low variation traits like Seed length and First flower (DAS) showed values from 4.76 to 7.15 mm and 31 to 51 days (Table 3). Comparing the distribution of across year-site BLUEs for the different traits, we found that for the traits Final height, Proportion of ripe fruits, Yield, Harvest (DAS), Lodging, Yield, TGW and Cover late, accessions showed near normal distribution. In contrast, the phenology traits First flower (DAS), First green fruits (DAS), First brown fruits (DAS) as well as Biomass showed distinct right-skewed distributions with most accessions clustered within the lower ∼25% of the range, whereas Early vigour and Cover were left-skewed (Figure 1A).

**Figure 1:**
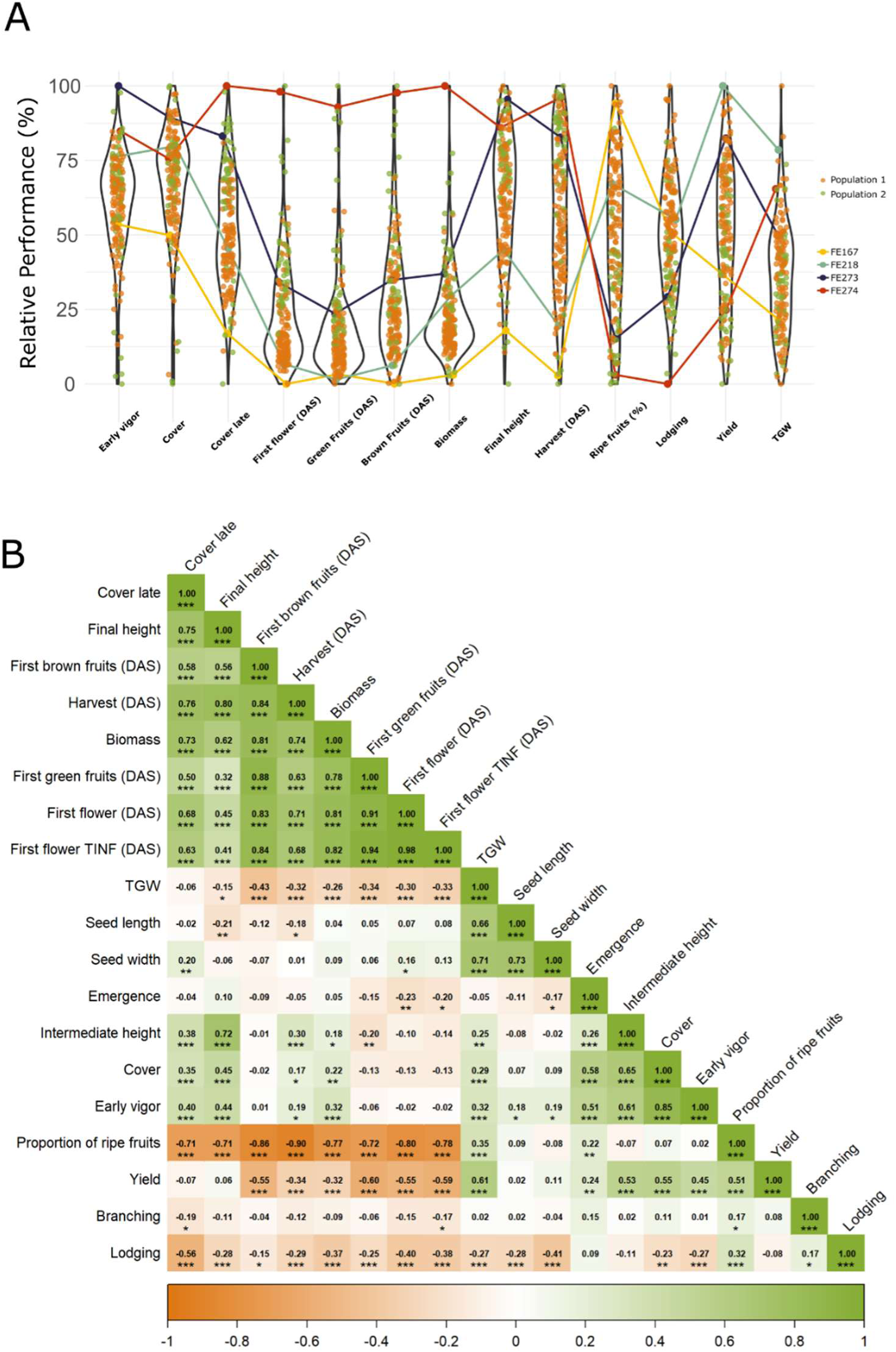
Distributions and correlations of across year-site best linear unbiased estimators (BLUEs) of the 1S agronomic traits investigated. **A** Points indicate the relative performance of all 170 accessions for the respective traits with colors indicating the accessions ancestral origin and violin plots indicating the distribution of accessions. Ancestral populations were assessed using NGSadmix with population 1 mainly representing accessions from Europe while population 2 focusing on Asia (Hess et al., 202Cb). The lines showcase the performance of four selected accessions, illustrating trait combinations within accessions with FE1C7 and FE218 being part of population 1 and FE273 and FE274 beloning to population 2. **B** Pairwise Pearson correlations (r) between BLUEs. Green coloration indicates positve, orange coloration negative correlations while saturation is proportional to their absolute correlation. Asterisks indicate the significance from two-sided correlation tests at different significance levels: ∗p < 0.05, ∗∗ p < 0.01 and ∗∗∗ p < 0.001.

**Table 3:** Summary statistics of best linear unbiased estimators (BLUEs) across year-sites. N_ys_, N_A_, DAS, CV and H^2^ refer to number of year-sites, number of accessions, days after sowing, coefficient of variation and heritability, respectively. The grades used for evaluating Branching, Early vigour and Lodging all range from 1 (low) to S (hight).

| Trait | Unit | $N_{ys}$ | $N_A$ | Mean | CV | Minimum | Maximum | $H^2$ |
| --- | --- | --- | --- | --- | --- | --- | --- | --- |
| First flower (DAS) | days | 4 | 170 | 35.64 | 10.16 | 30.81 | 51.09 | 0.97 |
| Seed width | mm | 3 | 170 | 4.09 | 10.27 | 3.01 | 5.41 | 0.97 |
| Seed length | mm | 3 | 170 | 5.82 | 6.19 | 4.76 | 7.15 | 0.95 |
| TGW | g | 2 | 166 | 23.49 | 16.73 | 15.68 | 37.32 | 0.95 |
| First flower TINF (DAS) | days | 3 | 170 | 41.69 | 12.23 | 35.56 | 63.69 | 0.94 |
| Yield | kg plot<sup>-1</sup> | 8 | 170 | 0.45 | 43.17 | 0.03 | 0.87 | 0.94 |
| Final height | cm | 7 | 170 | 115.6 | 13.07 | 68.65 | 142.05 | 0.93 |
| Harvest (DAS) | days | 4 | 170 | 120.04 | 14.83 | 79.23 | 149.18 | 0.93 |
| Proportion of ripe fruits | % | 2 | 163 | 52.55 | 46.15 | 5.79 | 100.89 | 0.93 |
| First green fruits (DAS) | days | 3 | 170 | 50.36 | 10.05 | 44.96 | 72.90 | 0.90 |
| Early vigor | grade | 4 | 170 | 5.60 | 13.93 | 2.90 | 7.35 | 0.86 |
| First brown fruits (DAS) | days | 3 | 170 | 72.95 | 11.73 | 60.21 | 104.12 | 0.85 |
| Cover | % | 6 | 170 | 50.92 | 12.94 | 26.76 | 62.31 | 0.84 |
| Biomass | kg plot<sup>-1</sup> | 2 | 169 | 2.23 | 36.32 | 0.88 | 6.54 | 0.82 |
| Cover late | % | 2 | 169 | 72.85 | 10.16 | 52.85 | 91.71 | 0.81 |
| Intermediate height | cm | 3 | 169 | 103.01 | 10.81 | 59.36 | 122.37 | 0.72 |
| Lodging | grade | 5 | 170 | 4.67 | 20.56 | 2.11 | 7.14 | 0.61 |
| Branching | grade | 2 | 164 | 7.33 | 8.59 | 2.85 | 8.32 | 0.51 |
| Emergence | count | 4 | 169 | 48.92 | 10.81 | 30.84 | 62.55 | 0.45 |

Looking at the performance of individual accessions for different traits revealed distinct agronomic profiles. In Figure 1A, the profiles of the four example accessions FE167, FE218, FE273 and FE274 are highlighted. Accessions FE167 and FE218 both showed a similarly fast early development (i.e., low values for First flower (DAS), First green fruits (DAS) and First brown fruits (DAS)), but FE218 had higher Early vigour, reached a greater Final height and had higher Yield. On the other hand, the accessions FE273 and FE274 were both harvested later and had a higher Final height compared to FE167 and FE218. However, FE273 had a faster early development and higher Yield compared to FE274.

### 4.2 Correlations between traits

Four trait groups were identified based on pairwise Pearson correlations among trait BLUEs across year-site combinations, while Branching and Lodging did not cluster with any group (Figure 1B). The first group consisted of eight traits related to late vegetative and generative stages of common buckwheat development (Cover late, Final height, Frist brown fruits (DAS), Harvest (DAS), Biomass, First green fruits (DAS), First flower (DAS), First flower TINF (DAS)) and showed correlation coefficients between each other ranging from 0.32 to 0.98 with a mean *r* of 0.71. The second group included traits describing seed shape (TGW, Seed length, Seed width) and had a mean *r* of 0.70. While Seed length and Seed width did not show a consistent correlation pattern with traits in other groups, TGW was positively correlated to Yield (*r* = 0.61) and Proportion of ripe fruits (*r* = 0.35) but negatively correlated to all trait of the first group (late development, mean *r* = - 0.27). The third group consisted of traits describing early vegetative development including Emergence, Intermediate height, Cover and Early vigour. These traits showed a within-group correlation coefficients between 0.26 and 0.85 (average r = 0.58). Taits in this group showed a consistent pattern of correlations (except Emergence) with traits from the first group (late development), as they were positively correlated to Cover late, Final height, Harvest (DAS) and Biomass but not significantly (*p* > 0.05) correlated to the other traits in group one. The fourth group includes only two traits (Proportion of ripe fruits and Yield), which were correlated with an *r* = 0.51. Both traits showed a consistent pattern of negative correlations with traits of group one. Branching did not show strong correlations with any other traits, while Lodging was negatively correlated to most traits of group one, two and three (-0.56 < *r* < 0.09) but was positively correlated to Proportion of ripe fruits.

### 4.3 Yield and yield stability across year-sites

Yield was highly correlated among the trials HOH24_early, HOH24_mid, LfL23, LfL24, and ETH24 (mean r = 0.93), based on Type B genetic correlations across year-site combinations. The year-sites JKI23 and JKI24_early were moderately correlated between each other (r = 0.8) and with the year-sites in the group mentioned above (0.68 < r < 0.91). In contrast, ETH23 showed low correlations (r < 0.6) with all year-sites except LfL23 (r = 0.73, Figure 2A). This pattern can also be observed in Figure 2B where all year-sites except ETH23 point approximately in the direction of the first principal component placing high yielding accessions on the left (e.g., FE218, FE059) and low yielding ones at the right (e.g., FE205, FE246). The second principal component was mainly influenced by ETH23 with accession performing especially well in this year-site such as FE181 and FE267 at the top of the plot. The factor analytic model for the multi-environmental data showed that factor 1 explained 38.12% of the variance between year-sites and separated the year-sites by placing ETH23 on one side and JKI23 and JKI24 at the other side of the scale (Figure 2C). Focusing on the relationship between Yield and Harvest (DAS), we found that in ETH23 accession with late harvest performed comparatively better than in other year-sites where the top yielding accession tended to show low Harvest (DAS) with LfL23 taking the middle ground were a few late accessions performed well but most high yielding accession were harvested relatively early (Figure 2C).Yield stability over year-sites expressed as Wricke’s ecovalence of all accessions are presented in Figure 3A and B. We found that accession with late harvest dates (Harvest (DAS) > 130) not only tended to have lower yields but these yields were also less stable. However, this pattern was not consistent when focusing on the five highest yielding accessions (FE218, FE059, FE212, FE281 and FE255) where the accession with the latest harvest (FE255) showed the highest stability (Figure 3A). Eight out of eleven accessions with determinate growth habit were among the 25% best yielding accessions but they also tended to have low yield stability compared to accessions of indeterminate growth habit with similar yields. Only one self-compatible accession (CC1003) fell within the 25% top yielding accessions, and it had a low to medium yield stability (Figure 3B).

**Figure 2:**
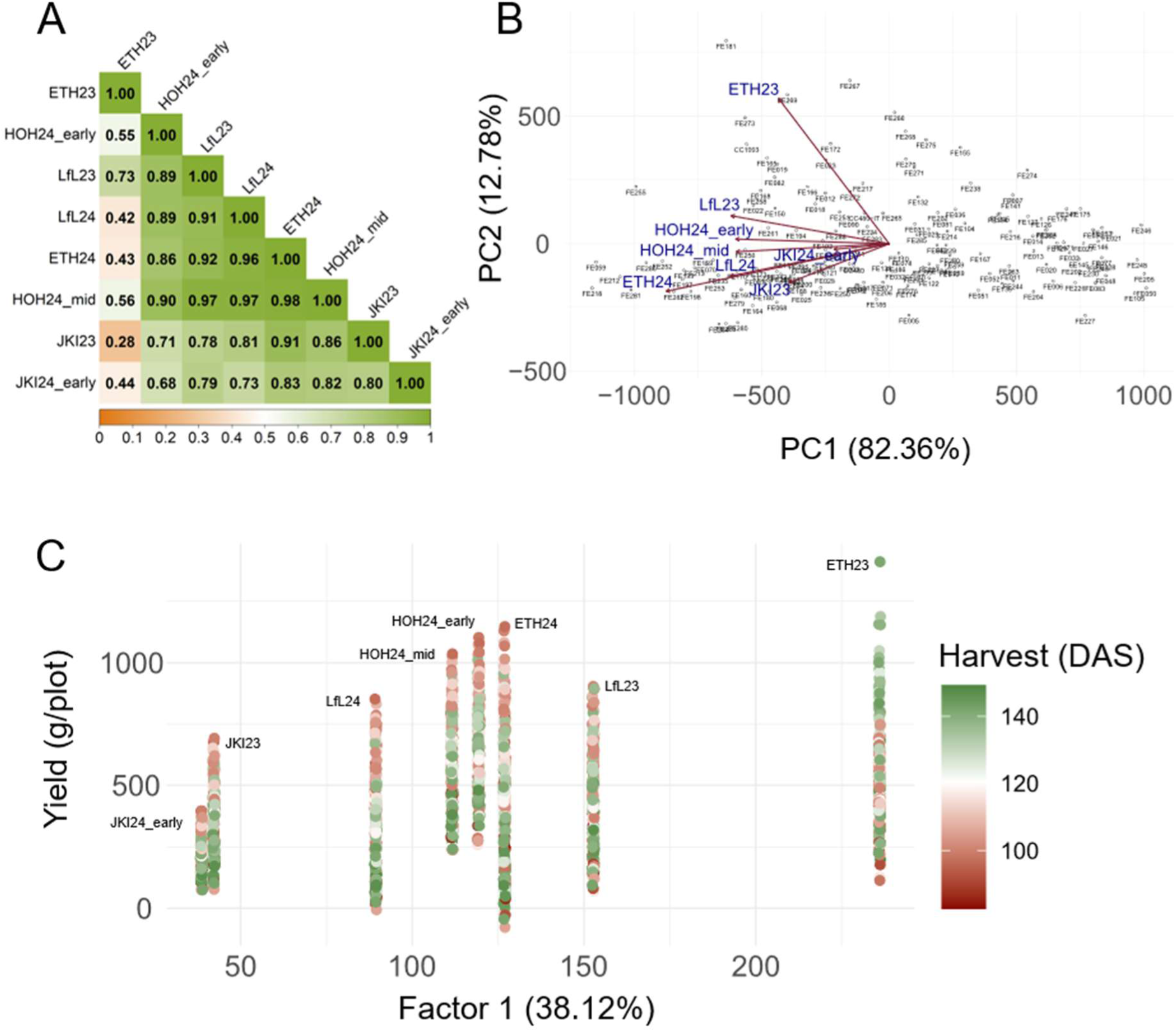
Yield stability across environments assessed by a factor analytic genotype-by-environment model (number of factors = 3): **A** Type B genetic correlations between yields (g/plot) of accessions grown in different year-sites. **B** Biplot showing the first two principal components based on yield of the 170 buckwheat accessions at all year-sites derived from the factor analytic model with red arrows indicating the effects of year-sites. **C** Predicted seed yields of all accessions across year-sites are displayed with dots being coloured according to the best linear unbiased estimator (BLUE, over all year-sites) of the harvest date (DAS). The x-axis represents the loading of factor 1 of the model for each year-site.

**Figure 3:**
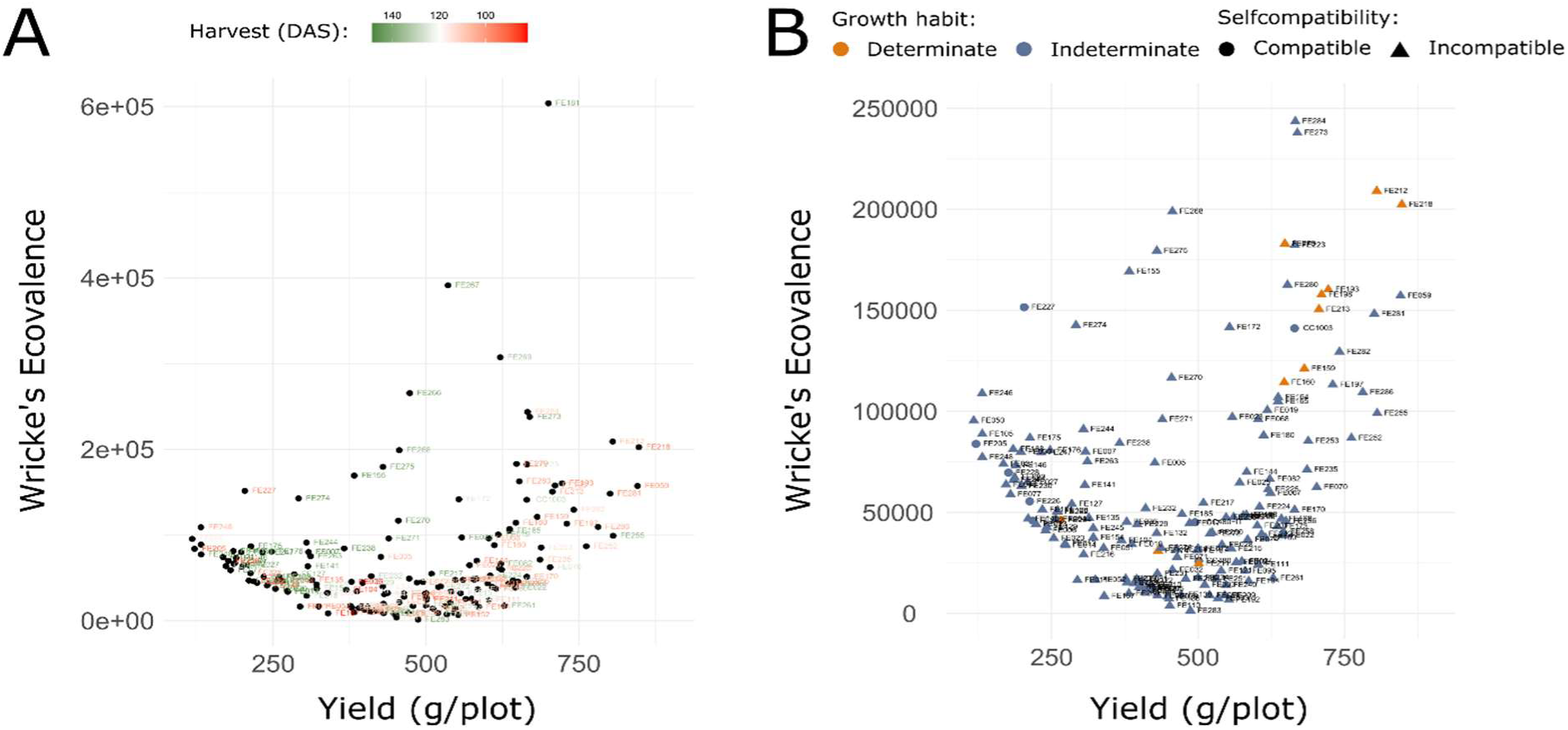
Yield stability across year-sites in relation to earliness and growth habit, shown as Wricke’s ecovalence. **A** Accession IDs are colored according to their harvest date (days after sowing, DAS) with red and green indicating early and late harvest dates, respectively. **B** Point color refers to growth habit and shape to self-compatibility. Note different y-scales in subplots: Four accessions (FE181, FE2C7, FE2CS and FE2CC) with the highest ecovalence scores were excluded in panel B for reasons of visibility. The three removed accessions all showed indeterminate growth habit and are self-incompatible.

### 4.4 Gene-trait associations

A total of 1,517 significant (p < 0.05) MTA for all assessed trait-year-site combinations were identified by combining the results from 328 GWAS performed on 82 trait-year-site combinations (Table 2) using four different models. Combining MTAs detected at multiple year-sites for the same traits results in 1,306 MTA for all traits with numbers of MTA per trait ranging from zero (Emergence) to 151 (Seed width). Finally, 1,116 unique SNPs were counted that showed a significant MTA with at least one trait (Supplementary Table S13, Supplementary Figure S2). Of these unique SNPs, 824 were localized within 471annotated genes, with 396 SNPs hitting a putative coding sequence (CDS) within these genes (Supplementary Table S15). Expanding the search radius around significant SNPs to 50 kb yielded a total of 3,681 putative genes associated with any of the assessed traits (Supplementary Table S14). As shown in Supplementary Figure S2, most MTA found for each trait only appear in one or few year-sites assessed. However, for the traits Biomass, First flower TINF (DAS), First green fruits (DAS) and Seed length, we observed at least one MTA that was significant in all year-sites as well as over year-sites. Moreover, 13 out of 14 traits showed at least one MTA that occurred in most of the year-sites where the trait was assessed (Supplementary Figure S2). Focusing on GWAS using across year-site BLUEs, we observed 22 SNPs that showed significant associations with two or more traits (Figure 4).

**Figure 4:**
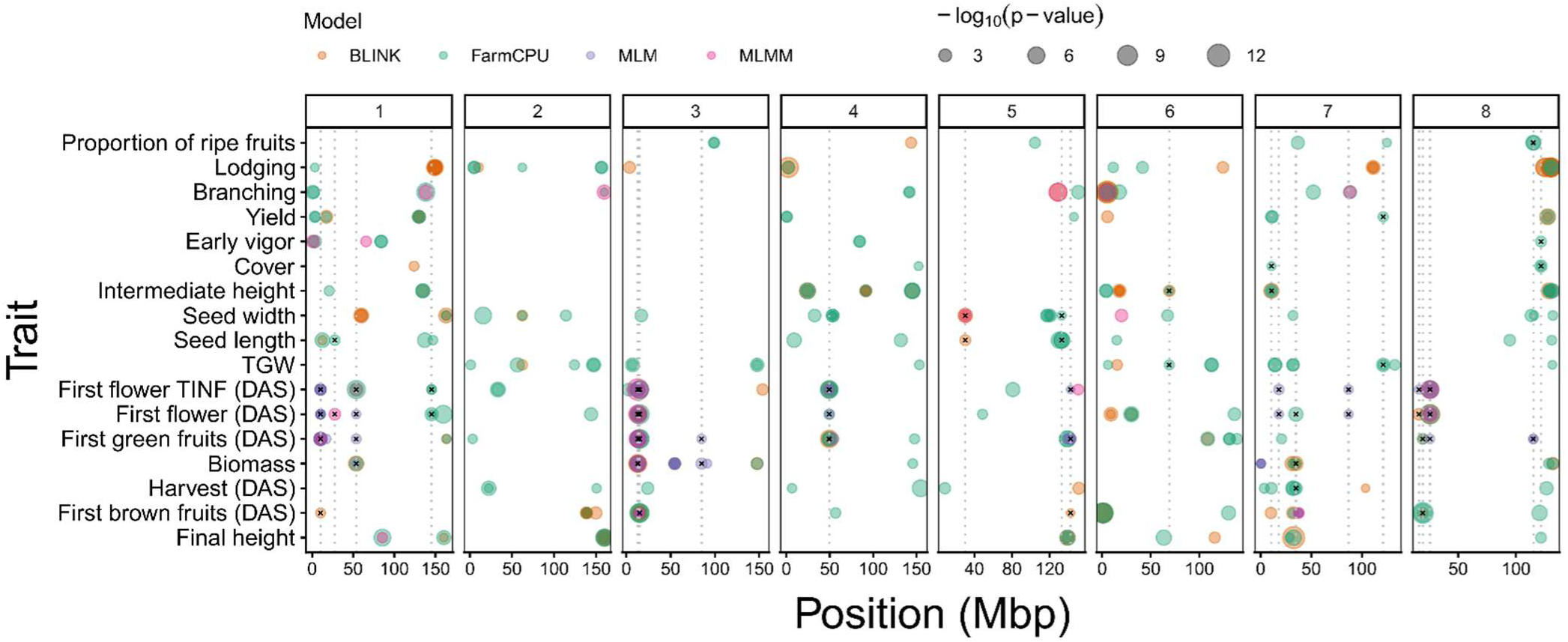
Results of the genome-wide association study (GWAS) based on best linear unbiased estimators (BLUEs) over all year-sites. Columns represent the eight chromosomes of buckwheat with positions within chromosomes indicated on the x-axis. The traits studied are listed on the y-axis. Each point represents a significant (p < 0.05) marker-trait association with colors indicating the model used (BLINK, FarmCPU, MLM, MLMM given in orange, green, violet and pink, respectively) and point size reflecting the Benjamini-Hochberg adjusted −log_10_(p) values. Dotted horizontal lines indicate the position of single nucleotide polymorphisms that show significant associations with two or more traits while black crosses mark the traits involved.

In Table 4, nine selected SNPs that were significantly associated with one or more traits are presented together with selected annotated genes found within 50 kb of these SNPs. The SNPs “chr1_53733173_G/A”, “chr3_12933889_G/C”, “chr4_49439763_G/A”, “chr8_115062905_C/T” were all associated with three or more BBCH stages. In addition, “chr1_53733173_G/A” and “chr3_12933889_G/C” were also significantly linked to Biomass while “chr8_115062905_C/T” was associated with Proportion of ripe fruits. Three of the presented SNPs (“chr5_132733474_A/G”, “chr5_30278292_T/C” and “chr6_19899220_C/T”) showed association with seed size related traits (Seed length and or Seed width). Finally, “chr7_120969696_T/C” was significantly associated with Yield and TGW while “chr8_121683930_A/C” showed a link to Cover and Early vigour. Except for “chr1_53733173_G/A” annotated genes were found in proximity (< 50 kb) for all presented SNPs.

**Table 4:** Selected marker-trait associations for single nucleotide polymorphisms (SNPs) that were significantly associated with multiple traits based on best linear unbiased predictors (BLUEs) across year-sites. SNP and gene ID refer to the reference genome Tuka while the annotation was adapted from the EggNOG (v.5.0.0) database (Huerta-Cepas et al., 201S). Distance to SNP indicates the distance between the significant SNP and the annotated gene in base pairs (BP).

| SNP | Traits<br>(over year-sites) | Gene ID | Annotation | Function | Reference | Distance to<br>SNP [bp] |
| --- | --- | --- | --- | --- | --- | --- |
| chr1_53733173_G/A | Biomass, First flower, First flower TINF, First green fruit | NA | NA | NA | NA | NA |
| chr3_12933889_G/C | Biomass, First flower, First flower TINF, First green fruits, First brown fruits | evm.TU.chr3.1088 | GRF1-interacting factor | Plant development and growth; photoperiod-sensitivity | (Lee *et al.*, 2009); (Piya *et al.*, 2020) | 0 |
|  |  | evm.TU.chr3.1087 | Auxin-induced protein 15A-like | Development of olive flowers; adaption of growth and development | (Salimonti *et al.*, 2021), (Stortenbeker and Bemer, 2019) | 6157 |
| chr4_49439763_G/A | First flower, First flower TINF, First green fruits | evm.TU.chr4.1245 | ARGONAUTE family | Effects on plant development and growth | (Zaheer *et al.*, 2024) | 0 |
| chr5_132733474_A/G | Seed length, Seed width | evm.TU.chr5.2258 | Sodium metabolite cotransporter BASS4 | Increase of seed size in Arabidopsis | (Lee *et al.*, 2017) | 0 |
| chr5_30278292_T/C | Seed width, Seed length | evm.TU.chr5.625 | Mitogen-activated protein kinase | Influence seed size; seed size in rice | (Zhang and Zhang, 2022), (Tian *et al.*, 2021) | 0 |
| chr7_120969696_T/C | Yield, TGW | evm.TU.chr7.3298 | MADS-box protein | Seed development, ripening and dehiscence, fruit and seed development | (Masiero *et al.*, 2011), (Adhikari and Kasahara, 2024) | 18367 |
| chr8_115062905_C/T | Proportion of ripe fruits, First flower | evm.TU.chr8.1963 | Catalyzes the ATP-dependent amination of UTP to CTP | Involved in endosperm development | (Yoon *et al.*, 2021) | 16462 |
|  |  | evm.TU.chr8.1964 | Cysteine-rich RLK (RECEPTOR-like protein kinase) 8 | Plant growth and development, delayed flowering | (Martin-Ramirez *et al.*, 2025) | 23220 |
| chr8_121683930_A/C | Cover, Early vigour | evm.TU.chr8.2333 | Multi antimicrobial extrusion (MATE) (TC 2.A.66.1) family | Regulator of plant growth and development, regulator of auxin levels in meristematic tissue | (Pastacaldi *et al.*, 2024) | 10925 |
| chr6_19899220_C/T | Seed width | evm.TU.chr6.1957 | F-box kelch-repeat protein | Differentially expressed during seed development, effects on seed color | (Jain *et al.*, 2007), (Hong *et al.*, 2020) | 44 |

## 5 Discussion

This study provides the most comprehensive catalogue of high-quality phenotypic data for common buckwheat to date, encompassing 19 agronomic traits evaluated across a large and diverse collection. The dataset is characterized by a high number of year-sites, as well as strong within year-site repeatabilities and across year-site heritabilities for most traits. Our GWAS results explore how high-quality data enables the discovery of MTA based on GWAFFs. Therefore, this resource provides a solid foundation for breeding improved varieties of common buckwheat that can fulfil various roles in the agricultural system.

### 5.1 Phenotypic diversity and the search for ideotype(s) of common buckwheat

To explore the idea of identifying ideotypes for common buckwheat, we selected four accessions with distinct trait combinations (Figure 1) that could be desirable depending on the crop’s role in the system. Based on this data, accession FE274 exhibits promising characteristics to function as a cover crop, as it shows good early establishment (Early vigour, Cover, Cover late), high Biomass production and low amount of viable seed (low Yield and low Proportion of ripe fruits) which are all considered favourable traits for a cover crop (Wilke and Snapp, 2008; McKenzie-Gopsill, 2025). The accessions FE218 and FE273 both match the requirements for a grain producing accession as they show good yields, but FE218 seems generally superior with higher Yield, earlier Harvest (DAS) and higher Proportion of ripe fruits. However, the better early development, higher Biomass production and lower Lodging of FE273 might make it an appealing option for organic agriculture, where weed competition and production of biomass to build up soil organic matter are especially important breeding goals beside increasing yield (Wolfe *et al*., 2008). Finally, FE167 shows generally weak performance for core agronomic traits (Early vigour, Lodging, Yield etc.). However, its early Harvest (DAS), low Biomass and high Proportion of ripe fruits might make it an interesting resource for breeding. These results show that phenotypic variation can be directly exploited to identify and select accessions for distinct breeding ideotypes.

### 5.2 Yield stability across environments

While high average seed yield is a primary breeding goal, yield stability under diverse environmental conditions is equally important (Calderini and Slafer, 1998). This is especially true for common buckwheat, as unstable yields are an often cited constraint of its productivity and economic viability (Jacquemart *et al*., 2012; Costanzo, 2019; Ohsawa, 2020). The analysis of yield stability (expressed as Wricke’s ecovalence) confirms the finding of Ohsawa (2020), that despite their generally high heritability, yields of common buckwheat tend to be relatively unstable between years. Our data suggest that accessions with average yield tend to be more stable while accessions with the highest (and lowest) yields show lower stability. In addition, accessions with late harvest tend to show lower yield stability compared to early accessions. The notable exception to this pattern is FE255, which has a relatively late harvesting time but shows a favourable combination of high and stable yields, that is only rivalled by the early accessions FE252 and FE286. Determinate growth habit and self-compatibility have both been described as target traits to breed for higher and more stable yields in common buckwheat (Matsui *et al*., 2007; Amelin *et al*., 2020). However, our results do not generally support the positive effect of these traits on yield stability. The high-yielding accessions with determinate growth habit showed comparatively low yield stability. Nevertheless, this trait remains a valuable breeding target as these accessions produce higher average yields across year-sites compared with traditional varieties. The relatively low and unstable yields of self-compatible accessions suggest that inbreeding depression in common buckwheat might not yet have been overcome, and further research and breeding efforts are required to harness potential productivity benefits of this trait.

The interpretation of yield stability estimations depends on the environments in which the data were collected. Type B genetic correlations of Yield between year-sites combined with the results of the PCA describing the year-site effects on Yield illustrate that results from ETH23 are distinct from other year-sites. Rainfall and temperature patterns during the growing season at ETH23 were characterized by a dry phase in mid-August that was ended by heavy rainfalls at the end of the month, coinciding with the ripening phase of many early accessions. In the same year the period from September to beginning of October (i.e., ripening period of late accessions) was generally dry and warm. Preharvest sprouting (PHS) in common buckwheat is usually severe if periods of humid and hot conditions occur right before harvest (Hara *et al*., 2007, 2020; Morishita *et al*., 2020; Suzuki *et al*., 2021; Takeshima *et al*., 2021). Very severe cases of PHS affecting the yield of early accessions such as FE213, FE218 and FE212 were observed in the field in ETH23 but could not be quantified. Late accessions did not show PHS as they were not ripening yet when the rainfall occurred and experienced favourable conditions (warm and dry) during their own ripening period. In LfL23, which was similar to ETH23 based on Type B genetic correlations, comparable but less pronounced weather patterns than in ETH23 were observed. In 2024 no comparable dry phases as in 2023 were recorded and rainy conditions dominated in September and October of 2024 at all seven year-sites, thus potentially explaining depressed yield for late accessions in 2024. This presumption is supported by yield data of the two check accessions FE213 (early harvesting) and FE273 (late) used in HOH24. These two accessions were the only ones that were included in all three harvesting groups and thus harvested at three different dates irrespective of their individual ripeness. While both accessions showed good yields when harvested with the early and mid-harvest group, yields dropped by two thirds for both the early and the late accession when harvested with the late harvest group. These observations illustrate that the large differences between accessions in terms of phenology and time of harvest require careful interpretation of the data, as accessions might experience different environmental conditions in specific developmental stages even when grown at the same year-site. Moreover, the data confirms that common buckwheat is highly sensitive to environmental conditions and changes in expected weather patterns might influence the choice of appropriate accessions.

### 5.3 Association genetics and the future of common buckwheat breeding

Different traits showed distinct patterns in terms of the number significant MTA detected, as well as the consistency with which these associations appeared at different year-sites. These patterns might be connected to different reactions of genotypes to environments or the complexity of the genetic control of these traits. The traits with lowest (Emergence) and highest (Seed width) heritability also yielded the lowest (0) and highest (151) number of MTAs, while across all traits the average correlation between heritability and the number of MTAs was r = 0.56. However, this relationship was not apparent for all traits. Traits related to BBCH stages showed high number of MATs regardless of their heritability whereas traits such as Early vigour or Proportion of ripe fruits showed high heritability but few MTA. This potentially reflects the high genetic complexity of traits related to phenological progress and flowering (Liu *et al*., 2021; Shi *et al*., 2022) compared to early vigour (Maulana *et al*., 2021), as reported for other plants. However, our results also identified a relatively small number of SNPs that showed significant associations with several traits and many of these associations were consistent over year-sites. Most of these associations involved traits that also showed high phenotypic correlations such as BBCH stages between themselves and with Biomass, Seed width with Seed length, Early vigour with Cover and Yield with TGW. As we only considered cases where the exact same SNP showed associations across traits and year-sites we might be underestimating the occurrence of such overlaps in cases where different SNPs in high linkage disequilibrium showed associations with multiple traits.

The identification of reliable MTAs that remain consistent in different environments under field conditions paves the way to targeted application of genomic methods in breeding of common buckwheat. Our results not only provide candidate MTAs that could be deployed for marker-assisted selection but also provide resources to implement genomic selection in common buckwheat. Examining the genome sequences around the respective MTA revealed biologically meaningful putative candidate genes underlying the respective traits, confirming both the validity of the GWAFFs genotyping method and its utility for GWAS. For example, SNP “chr3_12933889_G/C” which is associated with Biomass, First flower (DAS), First flower TINF (DAS), First green fruits (DAS) and First brown fruits (DAS) is located within a gene annotated as a GRF1-interacting factor. In Arabidopsis (*Arabidopsis thaliana*) this gene has been described to be involved in the control of plant growth and development as well as the regulation of photoperiod-sensitivity (Lee et al., 2009; Piya et al., 2020). Seed length and Seed width are both significantly associated with SNP “chr5_132733474_A/G” which co-localizes with an annotated gene coding for a Sodium metabolite cotransporter BASS4. This gene has been reported to influence seed size in Arabidopsis (Lee et al., 2017). Thus, our results provide candidate genes as potential targets for genome editing approaches which could be employed to confirm the causal relationship between the gene and trait or create improved accessions with improved flowering timing, biomass production or increased seed size.

### 5.4 Conclusion

The ETH Zurich/FOAG collection harbours great genetic diversity for several important agronomic traits. The detailed phenotypic description of the accessions combined with existing genomic resources represent a key resource to advance breeding of common buckwheat. Future work should focus on utilizing these resources to move towards the implementation of genomics-based approaches in applied common buckwheat breeding programs. Increased efficiency and speed of developing agronomically superior varieties of common buckwheat are necessary to boost its adoption in current and future agricultural systems. This will allow to harness the agronomic and ecological benefits of common buckwheat and ultimately contribute to broadening agricultural diversity.

## Authors contributions

**FH**: Writing – original draft, Conceptualization, Investigation, Formal analysis, Visualization, Data Curation. **LK**: Writing – review & editing, Conceptualization, Methodology, Software. **GR**: Writing - Review & Editing, Investigation. **HH**: Writing - Review & Editing, Resources, Project administration, Funding acquisition. **MHH**: Writing - Review & Editing, Project administration. **VB**: Writing - Review & Editing. **SY**: Writing – review & editing, Conceptualization, Investigation. **BS**: Writing – review & editing, Conceptualization, Methodology, Resources, Project administration, Funding acquisition.

## Acknowledgments

The authors would like to sincerely thank Simon Corrado for his invaluable help and expertise during the establishment, harvest and post-harvest processing of the field trials at Eschikon. We also thank Andreas Hund for his invaluable inputs in planning the field experiments and generous sharing of resources and expertise. We also acknowledge the valuable work of Shuyo Nodasaka, Benjamim Fabri, Manuel Cavicelli, Verena Knorst, Nico Helbig, Nicole Maier, Maximilian Mayr, Christine Kolm, Brigitte Pichlmaier, Andrea Deml, Alexandra Jestadt, Regina Achour and Dana Janowski supporting field work, collecting data and processing the harvest. Great thanks are due to Bianca Büttner for well-structured coordination of the project funded by the Federal Office for Agriculture and Food (BLE). Finally, we would also thank Carles Quesada Traver for providing scripts and inspiration for the analysis of the GWAS results.

## Data availability

Plot and accession level phenotyping data is available upon request.

