## Supplementary Figures for "Agronomic characterisation of a common buckwheat germplasm collection unlocks resources for genomics-assisted ideotype breeding": Supplementary_Figure_S1.pdf

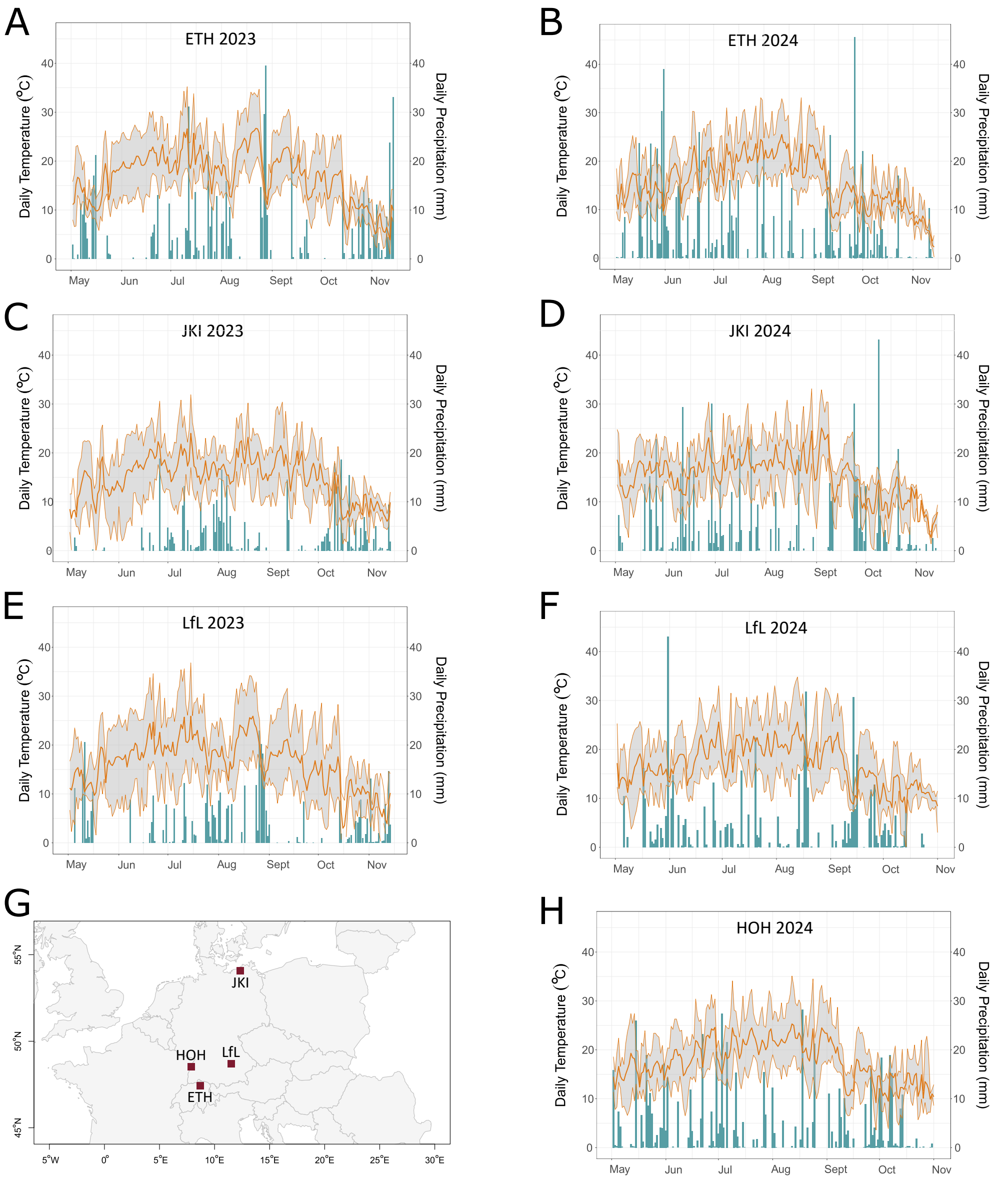


Supplementary Figure S1: Climate diagrams (A-F, H) and map locations (G) for the field trial sites. Thin orange lines indicate daily minimum and maximum temperatures while the fat line indicates daily temperature means. The blue bars indicate daily precipitation.
