## Supplementary Figures for "Agronomic characterisation of a common buckwheat germplasm collection unlocks resources for genomics-assisted ideotype breeding": Supplementary_Figure_S2.pdf

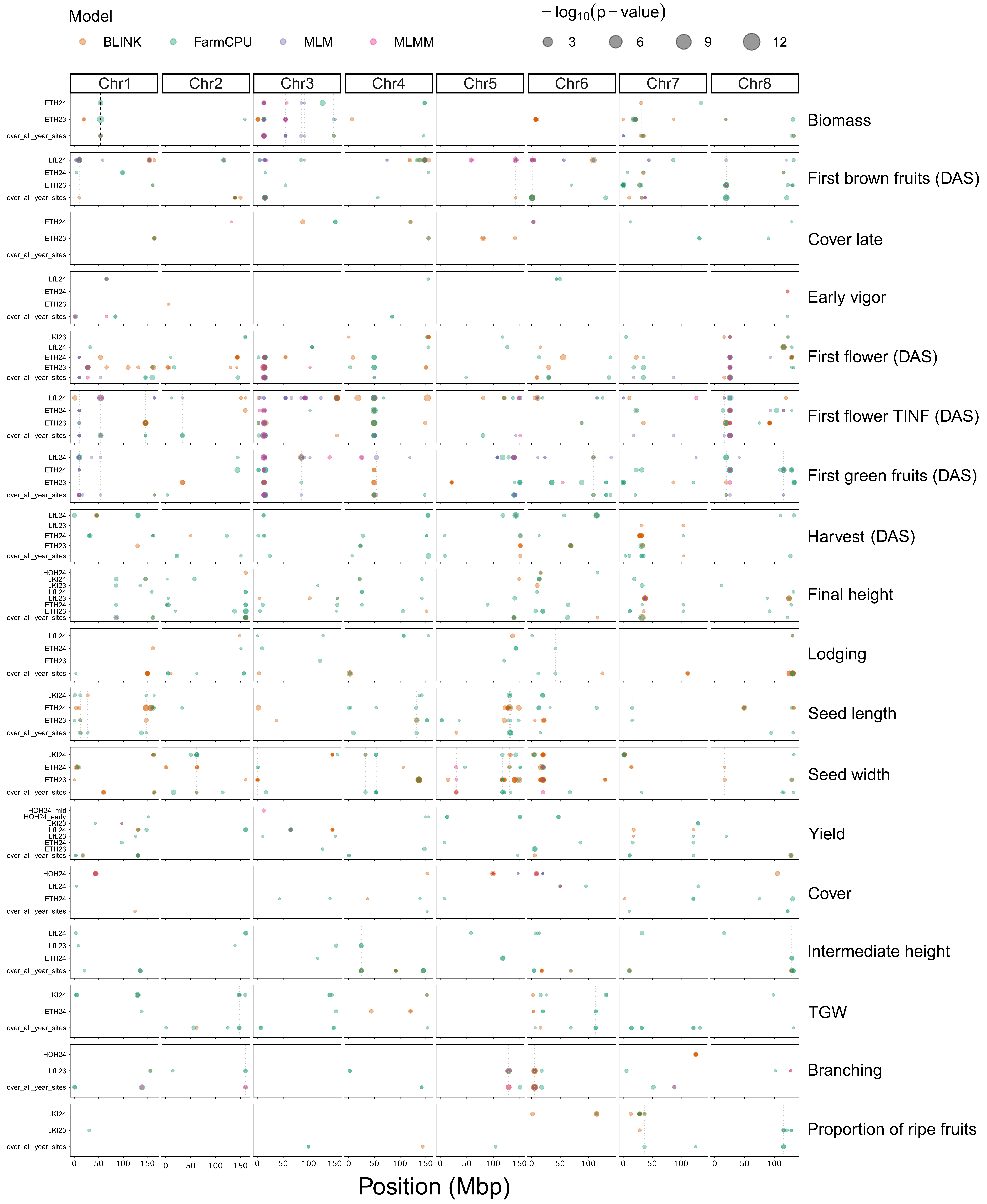


Supplementary Figure S2: Results of genome-wide association studies (GWAS) based on best linear unbiased estimators (BLUEs) of single year-sites as well as over all year-sites. Columns represent the eight chromosomes of the common buckwheat genotype Tuka (Hess et al., 2026a), while the studied traits are listed on the right-hand side y-axis. For each trait, the different year-sites where the trait was assessed and positive associations were detected are indicated on the left-hand side y-axis. Each point represents a significant ($p < 0.05$) marker-trait association with colors indicating the model used (BLINK, FarmCPU, MLM, MLMM given in orange, green, violet and pink, respectively) and point size reflecting Benjamini-Hochberg adjusted - 〖log〗 _10 (p) values (see legend). Within each trait panel, black dashed vertical lines show the positions of single nucleotide polymorphisms (SNPs) that yielded significant associations in all year-sites, while grey dotted lines refer to SNPs that showed associations in at least two year-sites and associations are missing in no more than two year-sites.
